# Single-nucleus transcriptional and chromatin accessibility profiling of mouse hypothalamic LepRb neurons reveals cell type-specific cis-regulatory elements linked to human obesity

**DOI:** 10.1101/2025.10.06.680592

**Authors:** Frankie D Heyward, Hui Pan, Jonathan M. Dreyfuss

## Abstract

Leptin receptor-expressing hypothalamic neurons (LepR^Hypo^) are key regulators of energy balance, yet a comprehensive, cell type-resolved, chromatin accessibility map of these neurons is lacking. We profiled ∼20,000 LepR^Hypo^ nuclei using single-nucleus multiome (snRNA-seq/snATAC-seq), identifying 39 transcriptionally and epigenetically distinct clusters, including AgRP (two subtypes), Pomc (two subtypes), Foxb1, Irx5/3 (three subtypes), Nts, PNOC (two subtypes), Kiss1/Pdyn (KNDy, two subtypes), Ghrh, Tcf7l2, and Sf1/Nr5a1 (three subtypes) populations. We also identified three Glp1r-expressing clusters with the highest Lepr enrichment, each marked by distinct molecular signatures. Cluster-specific open chromatin regions (OCRs) delineated putative cis-regulatory elements unique to each LepR^Hypo^ subtype. Mouse cell-type specific OCRs conserved in the human genome were identified; a subset were proximal to genes with high Human Genetic Evidence (HuGE) scores for obesity-related traits, overlapped obesity-associated GWAS loci, and/or coincided with eQTLs, including variants with the potential to influence human energy balance. Together, these data provide cell type-specific cis-regulatory atlas of LepR^Hypo^ neuronal subtypes, including Glp1r/Lepr-enriched populations, and highlight evolutionarily conserved, subtype-specific regulatory elements, associated candidate genes, and putative functional variants that may modulate LepR^Hypo^ subtype function and influence energy homeostasis and obesity susceptibility in humans.

## INTRODUCTION

The hormone leptin is a secreted adipokine that is a master regulator of energy homeostasis^1, 2^. During periods of weight gain, leptin is secreted from adipose tissue and binds to its receptor expressed on cell-types throughout the body, the lion’s share of which represent neuronal populations within the melanocortin circuit of the hypothalamus, and promotes physiological processes that ward against excess weight gain^2^.

Mice expressing a whole-body loss-of-function mutation in both Leptin or the Leptin Receptor become profoundly obese, and a phenotype that has also been exhibited by mice subjected to LepR deletion from AgRP neurons^3, 4, 5^. Given the indispensable role leptin-signaling within the hypothalamus plays in regulation of energy homeostasis, the hypothalamic cell-types upon which Leptin acts represent key targets for cell type-specific therapeutic intervention. A growing body of literature has identified transcriptional changes that occur within the broad family of LepR-expressing neurons using bulk-RNA-sequencing approaches, and AgRP neurons^6, 7, 8^.

More recently, efforts leveraging single-cell and single-nucleus RNA-seq (snRNA-seq) have been undertaken to profile the cellular diversity within individual hypothalamus subregions and whole hypothalamus, while identifying numerous energy-balance regulating neuronal types, including leptin-receptor expressing populations^9, 10, 11, 12, 13, 14, 15, 16^. Rupp et al. (2023), after performing snRNA-seq on LepR-expressing neuronal types, identified 18 clusters, including one that dually expressed both the highest degree of *Lepr* and *Glp1r*. Deletion of these LepRb^Glp1r^ neurons resulted in marked obesity and they were shown to mediate, in part, the weight-loss effects of semaglutide. More recently, Tan et al. (2024) performed snRNA-seq on all neuronal populations within the arcuate nucleus and identified 3481 cells, including AgRP and Pomc-neurons, as well as a novel *Bnc2*-expressing cell type that send inhibitory projections to AgRP neurons^17^. Furthermore, a recent paper used RAMPANT (Rabies Afferent Mapping by Poly-A Nuclear Transcriptomics) followed by snRNA-seq to identify various neuronal types that project onto AgRP neurons, and uncovered a LepR-enriched Trh/Cxcl12 population within the ARC that sends direct inhibitory projections onto AgRP neurons, and mediated the body weight-suppressing effects of the Glp1r agonist liraglutide^18^.

Open chromatin regions (OCRs) and their associated cis-regulatory elements (CREs) are principal regulators of transcriptional programs, and cell type-specific OCRs/CREs are predicted to encode unique cellular identity^19, 20^. We are still in the beginning stages of identifying putative cell-type CREs for rare neuronal types within the hypothalamus, let alone LepR-expressing subtypes. Inone et al., (2019) using bulk RNA-seq and ATAC-seq profiled stable CREs in the broad family of mouse LepR-expressing neurons^21^. Heyward et al. (2024), using AgRP neuron-specific RNA-seq and ATAC-seq, identified thousands of stable and dynamically regulated OCRs in response to fasting and leptin, while identifying putative CRE-binding transcription factors capable of coordinating transcriptional events during states of fasted-induced hunger and leptin-induced hunger suppression^7^. Advances in single-cell sequencing have enabled the ability to generate single-cell transcriptomes and chromatin accessibility profiles ^22, 23, 24^.

However, single-cell profiles of chromatin accessibility across the hypothalamus are newly emerging, with datasets existing across development (embryonic day 11 to postnatal day 8), and while comparing fed, a torpor-inducing 72-hour fasted, and refeeding for 12 hours or 1 week ^25, 26^. Yet, to date, no approach has employed a method for enriching for LepR-positive neuronal types prior to single-cell chromatin profiling, so as to appreciate the rich heterogeneity of CREs within underappreciated LepR subtypes, while linking them to transcriptional regulatory programs that influence energy homeostasis.

Here we generate single-nucleus RNA-seq (snRNA-seq) and ATAC-seq (snATAC-seq) profiles from mouse LepR-expressing cell-types within the mouse hypothalamus (LepR^Hypo^). We identify 39 different LepR^Hypo^ clusters, including those enriched for AgRP (two subtypes), Pomc (two subtypes), Bnc2/Nkx2-4/Glp1r (three subtypes), Foxb1, Irx5/3 (three subtypes), Nts, PNOC (two subtypes), Kiss1/Pdyn (KNDy, two subtypes), Ghrh, Tcf7l2, and Sf1/Nr5a1 (three subtypes), among others. We next identified cell-type specific CREs, and link each of these cell-type specific CREs to human coding and non-coding genomic loci associated with obesity.

## RESULTS

As a means of enriching for LepR-expressing cell-types within the hypothalamus, we crossed LepR-ires-cre mice to NuTRAP mice, thereby generating NuTRAP^LepR^ mice that enable the cre-dependent tagging of nuclei with eGFP ^27, 28^. 120,000 Nuclei from 6 female pooled hypothalami were collected via Fluorescence fluorescence-activated nuclei sorting (FANS), and subjected to Chromium Next GEM Single Cell Multiome ATAC + Gene Expression, with 4 libraries being made in parallel, with each library comprised of a different set of ∼5,000 nuclei (**Fig. 1a, Fig. S1a**). Across our 22,581 nuclei, we detected 39 different cell clusters, each with a distinct assortment of molecular identifiers and varying numbers of nuclei according to our snRNA-seq results (**Fig. 1a-c**).

**Figure 1.**
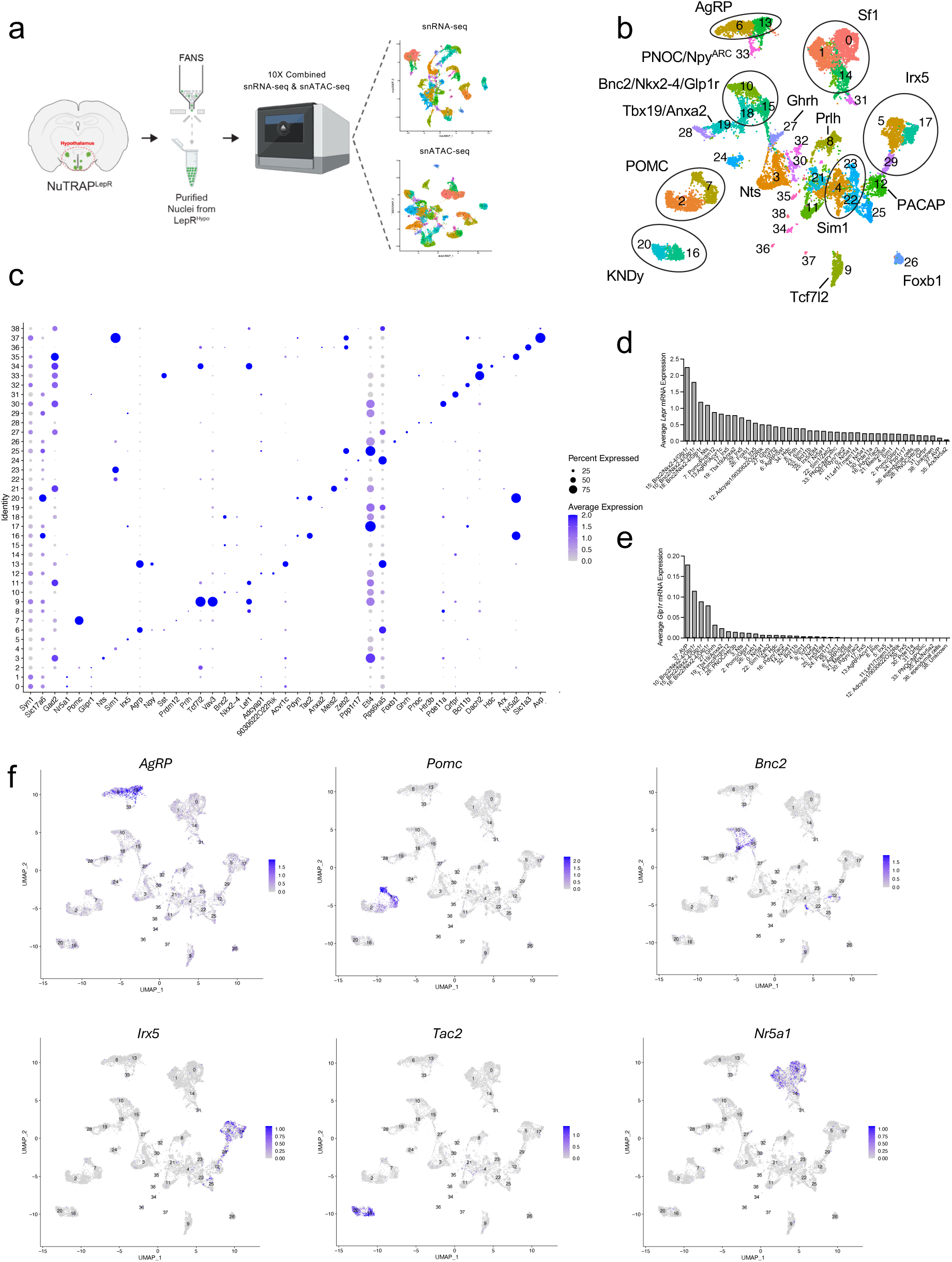
snRNA-seq profiles of 39 LepR^Hypo^ populations. (A) Schematic showing LepR^Hypo^ single-nucleus multiome profiling workflow. (B) snRNA-seq UMAP plot revealing 39 distinct populations. (C) dot plot showing cluster-specific transcriptional profiles. (D) Bar chart showing cluster-specific *Lepr* expression levels. (E) Bar chart showing cluster-specific *Glp1r* expression levels. (F) UMAP plots showing particular enriched genes (Agrp, Pomc, Bnc2, Irx5, Tac2, Nr5a1).

### Revealing the rich cellular heterogeneity within our LepR^Hypo^ snRNA-seq dataset

We detected 3 clusters expressing nuclear receptor subfamily 5 group A member 1 (*Nr5a1*), which is also called Steroidogenic factor-1 (SF-1) (Cluster 0: *Nr5a1*, Cluster 1: *Nr5a1*, and Cluster 14: *Nr5a1*)^29, 30^. These neurons are known to be expressed within the ventromedial hypothalamus (VMH). Clusters 2 and 7 comprise Pomc-expressing neurons within the ARC ^31^. Cluster 7 expresses ∼4 times more LepR than Cluster 2, and *Prdm12*, whereas Cluster 2 expresses *Glipr1*, a marker that is indicative of a recently characterized population of low LepR-expressing Pomc neurons (Cluster 2: *Pomc*/*Glipr1*, Cluster 7: *Pomc*/*Prdm12*) ^32^. Cluster 3 represents *Nts*-expressing neurons found in both the LH (Cluster 3: *Nts*) ^33, 34, 35^. Clusters 4, 22, and 23 represent *Sim1* neurons found in the PVH (CITE) (Cluster 4: *Sim1*, Cluster 22: *Sim1*, Cluster 23: *Sim1*). Clusters 5, 17, and 29 are *Irx5/Irx3*-expressing neurons of the ventral premammillary nucleus (Cluster 5: *Irx5*, Cluster 17: *Irx5*, Cluster 29: *Irx5*) ^36, 37^. Clusters 6 and 13 represent AgRP-expressing neurons found within the ARC. Cluster 6 is relatively enriched for *Sst*, whereas we observed that Cluster 13 is uniquely enriched for *Acvr1c* (Cluster 6: *Agrp*/*Sst*, Cluster 13: *Ag*rp/*Acvr1c*) ^38^. Cluster 8 comprise *Prlh-*expressing neurons found within the DMH (Cluster 8: *Prlh*)^39^. Cluster 9 represents *Tcf7l2*-expressing neurons of the DMH and Lateral Hypothalamus (LH) (Cluster 9: *Tcf7l2*) ^40^. Clusters 10, 15, and 18 represent the class of Glp1r-expressing neurons that also exhibit the highest *Lepr-*expression, a characteristic they share in common with a previously identified population of LepR^Glp1r^ neurons (**Fig. 1d**) ^39^.

Moreover, Clusters 10, 15, and 18 also exclusively express *Bnc2*, a newly identified marker for a Lepr-expressing and leptin-activated neuronal population that sends inhibitory projections to AgRP neurons ^17^. Furthermore, these neurons are greatly enriched for *Nkx2-4*, a previously identified marker of hypothalamic neurons ^41^. Thus, we henceforth refer to these neurons as *Bnc2*/*Nkx2-4*/*Glp1r* neurons. Cluster 11 represents a neuronal population expressing *Lef1* and *Tmem114* (Cluster 11:Lef1/Tmem114), which may be the mouse equivalent of a Lef1/Tmem114-expressing population recently identified within the human hypothalamus ^42^.

Cluster 12 expresses the highest amount of *Adcyap1*, a marker for PACAP neurons, along with *9030622O22Rik* (Cluster 12: *Adcyap1/*9030622O22Rik) and potentially represent previously characterized PACAP^PVN^ neurons ^43^. Clusters 16 and 20 represent neurons co-expressing Kisspeptin, neurokinin B, and dynorphin (KNDy) that reside in the ARC (Clusters 16: *Pdyn*/*Tac2*, Cluster 20: *Pdyn*/*Tac2*)^44^. Cluster 19 are Tbx19-expressing neurons, a population of previously identified neurons expressed in the ARC and have been previously show to be enriched for *Pirt* and we observed as enriched for *Anxa2* (Cluster 19: *Tbx19/Anxa2*)^18, 39^. Cluster 21 appear to be a previously identified population of *Meis2*/*Sst* expressing neurons that reside in the lateral hypothalamus area (LHA) (Cluster 21: *Meis2*/*Sst*)^14^. Cluster 24, are *Ppp1r17*-expressing neurons of the DMH (Cluster 24: *Ppp1r17*) ^45^. Cluster 25 expresses *Irx5* but no *Irx3*, while being greatly enriched for the gene Epilepsy, Familial Temporal Lobe, 4 (*Elt4*), thus we labeled them *Irx5*/*Elt4* neurons (Cluster 25: *Irx5*/*Elt4*). Cluster 26 are previously identified *Foxb1*-positive neurons, that potentially reside in the ventral medial hypothalamus (Cluster 26: *Foxb1*)^39, 46^.

Cluster 27 are Ghrh-positive neurons within the ARC (Cluster 27: *Ghrh*) ^39, 47, 48^. Cluster 28 are *Pnoc*- and *Htr3b*-positive neurons within the ARC (Cluster 28: *Pnoc*/*Htr3b*)^41, 49^. Cluster 30 appear to be a previously unidentified population of neurons that are enriched for Phosphodiesterase 11A (*Pde11a*) (Cluster 30: *Pde11a*). Cluster 31 are a previously identified population of *Qrfpr-*expressing neurons within the VMH (Cluster 31: *Qrfpr*)^50^. Cluster 32 appear to be a previously unidentified population of neurons that are enriched for *Bcl11b* (Cluster 32: *Bcl11b*). Cluster 33 expressed the most *Sst* compared to any other cluster, long with considerable *Otp* and *Npy*. Interestingly, Cluster 33 did not express *Agrp* despite abutting Clusters 6 and 13 in UMAP space. With this population also expressing *Pnoc*, but not *Htr3b*, as well as *Npy* and an exceptionally high amount of Sst, we believe these are the recently profiled PNOC/NPY^ARC^ neurons (Cluster 33: PNOC/NPY^ARC^) ^51^ (**Fig. S2a**). Cluster 34 represent histidine decarboxylase-expressing neurons (*Hdc*)-expressing neurons of the Tuberomammillary nucleus (Cluster 34: *Hdc*)^52^. Cluster 35 are possibly previously identified *Arx*/*Nr5a2* neurons given the enriched expression of those transcripts (Cluster 35: *Arx*/*Nr5a2*) ^10^. Cluster 36 are suspected ependymal cells given their expression of *Slc1a3*, and *Cd34* (Cluster 36: ependymal cells).

Cluster 37 are Arginine vasopressin (AVP)-positive neurons found in the paraventricular nucleus of the hypothalamus (PVH) (Cluster 37: AVP)^53^, given their expressing *Sim1*. Interestingly, these neurons appear to be greatly enriched for Glp1r (**Fig. 1e**). Cluster 38 did not present with a notable marker gene, and thus are classified as unknown (Cluster 38: Unknown).

Given the expression of Vesicular Glutamate Transporter 2 (*Vglut2*), alternatively called Solute carrier family 17 member 6 (Slc17a6), the following clusters were deemed to be glutamatergic: Clusters 0, 1, 5, 9, 12, 14, 16, 17, 20, 22, 23, 24, 25, 26, 29, 31, 37 (**Fig. S1c**). Meanwhile, given the expression of Glutamate Decarboxylase 1 (Gad1) and/or Gad2, which is required for the production of gamma-aminobutyric acid (GABA), we believe the following clusters correspond to GABAergic neurons: 3, 6, 10, 11, 13, 15, 18, 19, 21, 27, 28, 30, 32, 33, 34, 35, 38 (**Fig. S1c**). Cluster 36 does not express either canonical glutamatergic or GABAergic markers, nor does it express the general neuronal marker Synapsin I (*Syn1*), thus we suspect that it represents an non-neuronal LepR-expressing cell type. Pomc-neuron clusters 2 and 7 express appreciable amounts of Gad1/2, and more modest amounts of Slc17a6, perhaps reflecting the heterogeneity within this population, with 40% of POMC neurons being GABAergic^54^. Similarly, Clusters 4 and 8 also appeared to be dually glutamatergic and GABAergic given their expression profiles (**Fig. S1c**).

We performed label transfer to compare our snRNA-seq profiles with those from HypoMap, an extensive single-cell/nucleus atlas of murine hypothalamus cell types generated by Steuernagel & Lam et al. (2022)^41^. 19 of our Clusters were associated with cell types identified via HypoMap, suggesting broad concordance between our dataset and established hypothalamic cell type annotations (**Fig. S1d**).

### Unique molecular identifiers of AgRP and Bnc2/Nkx2-4/Glp1r neuronal subtypes

In our snRNA-seq data, we identified unique molecular identifiers that can be used to distinguish Cluster 6 and Cluster 13 AgRP neurons. Cluster 6 is enriched for *Sst*, *zfp804a*, and *cntn5*, while Cluster 13 expresses activin A receptor type 1C (Acvr1c) also known as Activin receptor-like kinase 7 (Alk7). Spatial transcriptomics using Xenium revealed that *Sst* transcripts were greatly enriched in ∼42% *Agrp* transcript-positive neurons, *Acvr1c* transcripts were identified within ∼42% of *Agrp* transcript-positive neurons, ∼10% of AgRP neurons expressed both Sst and Acvr1c, and ∼7% expressed neither transcript, all within the ARC (**Fig. 2a-d**).

**Figure 2.**
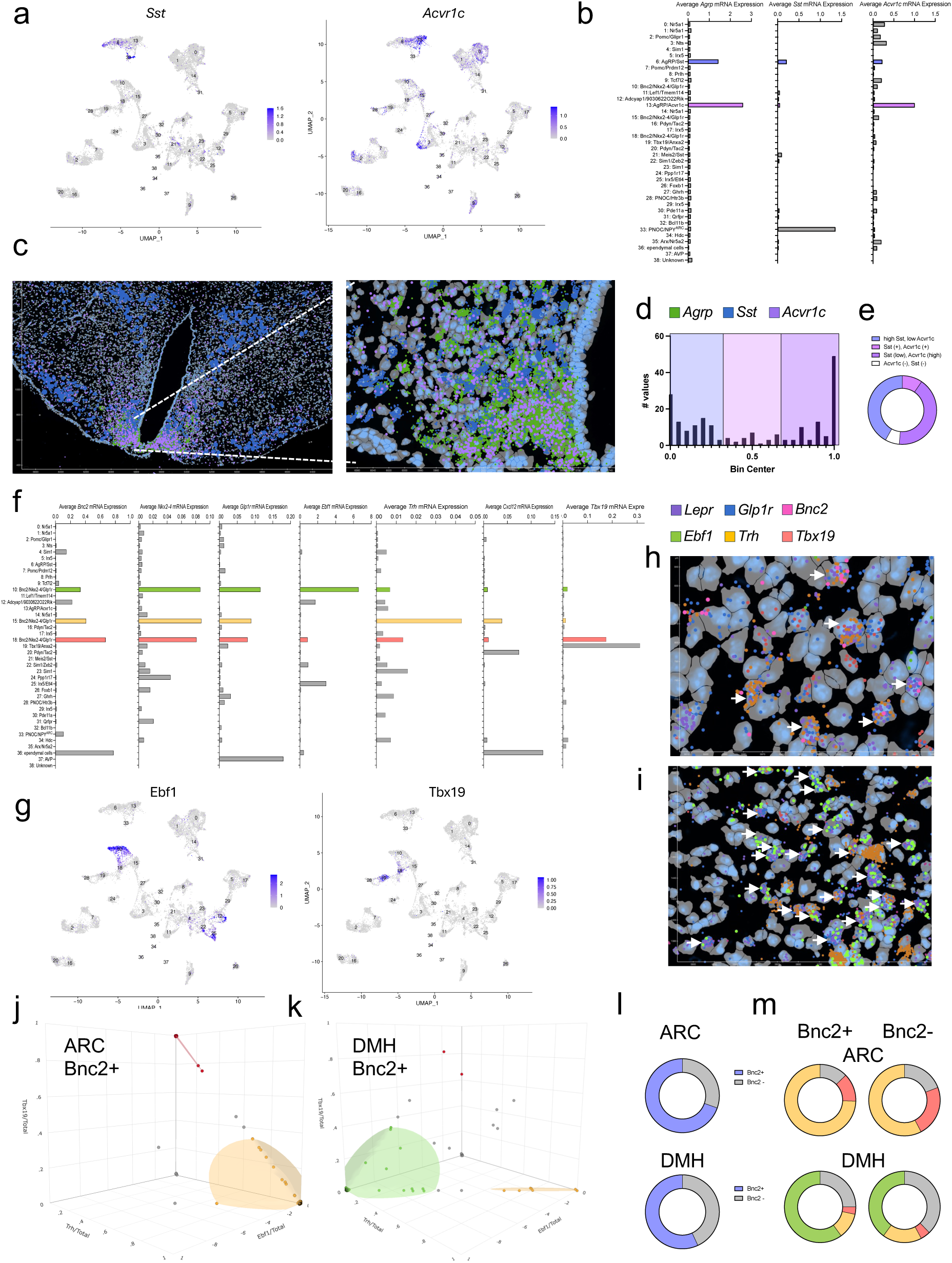
Spatial transcriptomic profiling of AgRP and Bnc2/Nkx2-4/Glp1r subpopulations. (A) UMAP plots for *Sst* (left) and *Acvr1c* (right) expression. (B) Bar graphs showing cluster specific Sst (left) and Acvr1c (right) expression. (C) Xenium explorer image showing AgRP (green), Sst (blue), and Acvr1c (lavender). (D) Histogram showing the binned count of cells with a particular degree of *Acvr1c transcript count /*(*Acvr1c + Sst transcript count*). (E) Doughnut chart showing the proportion of different neuronal populations. (F) Bar graphs showing the cluster-specific expression of, from leftmost to rightmost, *Bnc2*, *Nkx2-4*, *Glp1r*, *Ebf1*, *Trh*, *Cxcl12*, or *Tbx19*. (G) UMAP plot showing *Ebf1* (left) and *Tbx19* (right) expression. Xenium Explorer image within the (H) ARC/VMH and (I) DMH showing transcripts for *Lepr, Glp1r, Bnc2, Ebf1, Trh, Tbx19*. 3D Scatter Plots showing Ebf1/Total, Trh/Total, and Tbx19/Total within the (J) ARC//VMH (left) and (K) DMH (right). (L) Doughnut plots showing the proportion of Lepr and Glp1r co-expressing cells that are Bnc2-positive or Bnc2-negative. (m) Doughnut plots showing the proportion of Lepr and Glp1r co-expressing cells that are Ebf1+, Trh+, Tbx19+, or neither, while being either Bnc2-positive or Bnc2-negative.

We observed from our snRNA-seq data that there were three *Bnc2*/*Nkx2-4*/*Glp1r* neurons-positive Clusters (10, 15, 18) that each have unique molecular identifiers. Given the lack of published evidence of distinct *Bnc2*/*Nkx2-4*/*Glp1r* populations, we sought to further characterize this family of neurons. While all three express the most *Lepr* compared to the others, Cluster 15 has the most, followed by Cluster 10, and then Cluster 18 (**Fig. 2f**). Moreover, Cluster 10 is greatly enriched for the transcription factor *Ebf1*, compared to all other clusters (**Fig. 2f, g**).

Cluster 15 is enriched for *Trh* and *Cxcl12* (**Fig. 2f, S1b**). Cluster 18 is the only of the three to express the transcriptional regulator *Tbx19.* (**Fig. 2f, g**). We then employed spatial transcriptomics using Xenium to independently confirm the existence of these three *Bnc2*/*Nkx2-4*/*Glp1r* neuronal subclusters. We first confirmed there to be a considerable number of *Lepr* and *Glp1r* co-expressing neurons within the ARC/VMH and DMH (**Fig. 2h, i**). We observed that, within the ARC/VMH, ∼68% of *Lepr*/*Glp1r*-co-expressing neurons also express *Bnc2*, whereas ∼32% of *Lepr*/*Glp1r*-co-expressing neurons did not (**Fig. 2l**), suggesting that Bnc2 does label the majority of *Lepr*/*Glp1r*-co-expressing neurons in the ARC/VMH. Furthermore, 74.4% of *Lepr*/*Glp1r*/*Bnc2*+ neurons were relatively enriched for *Trh*, ∼13% for *Tbx19*, and 0% for *Ebf1* (**Fig. 2h, j, m**). Amongst *Lepr*/*Glp1r*/*Bnc2*-neurons, ∼58% were relatively enriched for *Trh*, ∼23% for *Tbx19*, and 0% for *Ebf1* (**Fig. 2h, j, m**). Thus, we concluded that the majority of *Lepr*/*Glp1r*/*Bnc2*+ and *Bnc2-* neurons within the ARC/VMH are enriched for Trh, with ∼13-23% being enriched for *Tbx19*, confirming the identity of two LepR/Glp1r neuron subpopulations within ARC/VMH.

We next confirmed that, within the DMH, ∼58% of *Lepr*/*Glp1r*-co-expressing neurons also express Bnc2, whereas ∼43% of *Lepr*/*Glp1r*-co-expressing neurons did not (**Fig. 2l**), suggesting that while Bnc2 does label the majority of *Lepr*/*Glp1r*-co-expressing neurons in the DMH, a substantial proportion of *Lepr*/*Glp1r*-co-expressing neurons do not appear to express *Bnc2*.

Furthermore, ∼12% of *Lepr*/*Glp1r*/*Bnc2*+ neurons were relatively enriched for *Trh*, ∼3% for *Tbx19*, ∼60% for *Ebf1*, and 25% were not enriched for either transcript (**Fig. 2i, k, m**). Amongst *Lepr*/*Glp1r*/*Bnc2*-neurons, ∼18% were relatively enriched for *Trh*, ∼4% for *Tbx19*, ∼40% for *Ebf1*, and 38% were not enriched for either transcript (**Fig. 2i, k, m**). Thus, we concluded that the majority of *Lepr*/*Glp1r*/*Bnc2*+ and *Bnc2-* neurons within the DMH are enriched for Ebf1, with ∼12-18% being enriched for *Trh*, and 3-4% enriched for Tbx19, confirming the identities of three LepR/Glp1r neuron subpopulations within the DMH (**Fig. 2i, k, m**).

### Identifying cell type-specific chromatin accessibility profiles across LepR^Hypo^ subtypes

We next sought to survey the chromatin accessibility landscape of LepR^Hypo^ neurons during basal conditions. As part of our Chromium Next GEM Single Cell Multiome ATAC + Gene Expression workflow in parallel with our snRNA-seq library preparation, we generated single-nucleus Assay for Transposase-Accessible Chromatin using-sequencing (snATAC-seq) libraries from the same nuclei, with our seeing 39 distinct clusters (**Fig. 3a**). We detected between 51011 and 129838 peaks in each of our clusters (**Fig. 3b**).

**Figure 3.**
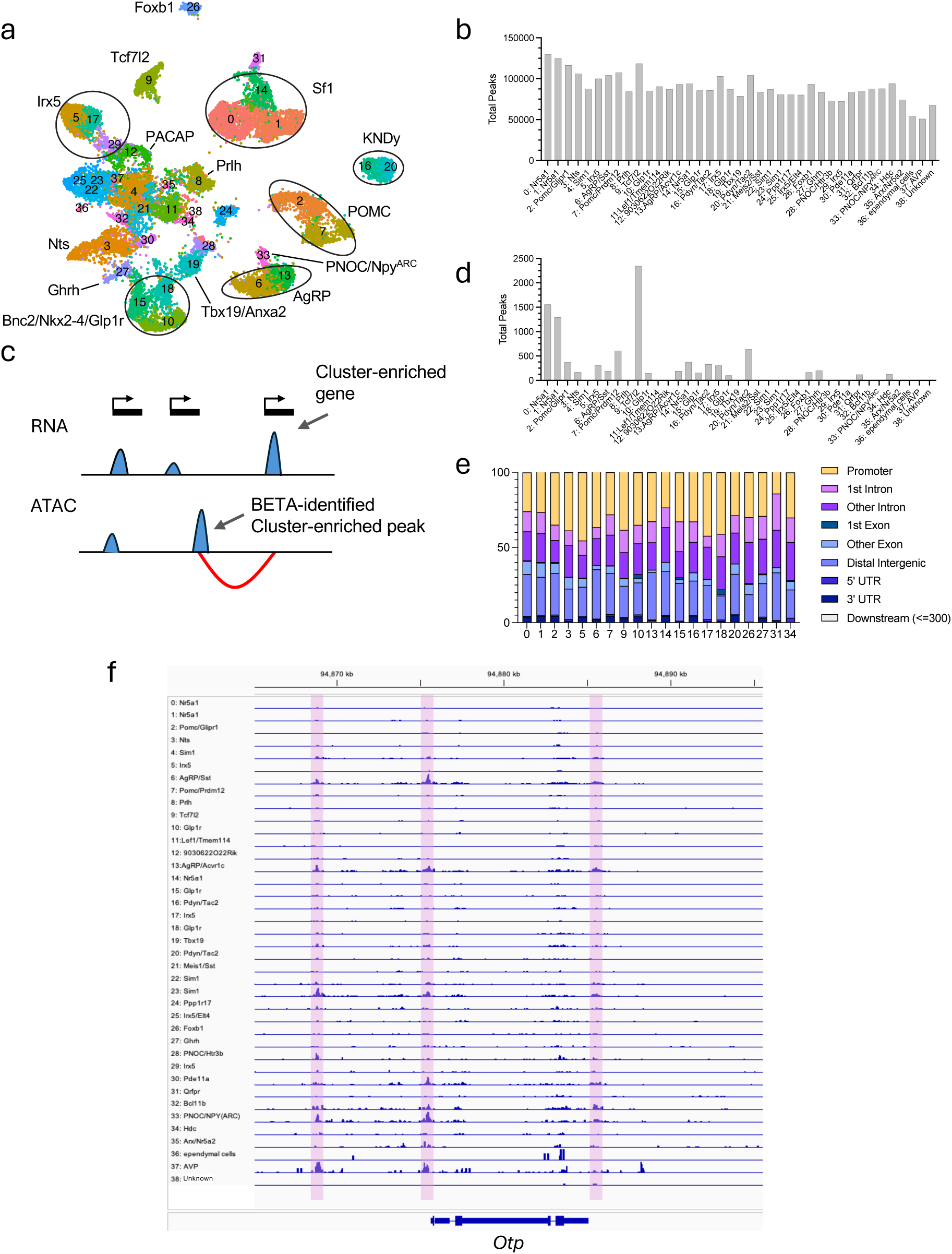
snATAC-seq profiles of LepR^Hypo^ populations. (A) snRNA-seq UMAP plot revealing 39 distinct populations. (B) Bar chart showing the number of open chromatin regions (OCRs) across each of the clusters. (C) Crude schematic showing the relationship between BETA-peaks and associated genes. (D) Number of BETA peaks in select clusters. (E) Parts of whole bar graph showing the proportion of BETA peaks found in different genomic elements. (F) IGV image showing pseudobulk snATAC-seq peaks within the vicinity of the Otp gene.

We next identified OCRs that were specific to either individual clusters or to pre-specified cluster families: AgRP (clusters 6: Agrp/Sst and 13: Agrp/Acvr1c), POMC (clusters 2: Pomc/Glipr1 and 7: Pomc/Prdm12), LepR/Glp1r (clusters 10: Bnc2/Nkx2-4/Glp1r/Ebf1, 15: Bnc2/Nkx2-4/Glp1r/Trh, and 18: Bnc2/Nkx2-4/Glp1r/Tbx19), and Kiss1 (clusters 16: Kiss1 and 20: Kiss1/Ak5), and within ±1 Mb of an annotated gene.

To assess whether cell-type-specific chromatin accessibility patterns are predictive of gene expression, we applied Binding and Expression Target Analysis (BETA) to each cluster using the corresponding differentially accessible ATAC-seq peaks and differentially expressed RNA-seq markers ^55^. Cluster-specific peaks were defined using the Wilcoxon Rank Sum test (adjusted p < 0.05), and cluster marker genes were similarly identified with Seurat. BETA calculates a regulatory potential (RP) score for each gene by integrating both the proximity and the enrichment of nearby peaks within ±100 kb of transcription start sites. For many clusters, including clusters 0, 1, 2, 3, 5, 6, 7, 9, 10, 13, 14, 15, 16, 17, 18, 20, 26, 27, 31, and 34, the upregulated genes exhibited significantly higher RP scores compared to both downregulated and non-significant genes (Kolmogorov-Smirnov test, p < 0.05, **Fig. S3a**). For these 20 clusters we observed between 100-2,344 OCRs with significant BETA scores (**Fig. 3d**). The majority of BETA peaks OCRs were found within intergenic (36-45%) and intronic (43–50%) regions, while only 3-11% of OCRs were found within gene promoter regions, defined as regions <1000 bps upstream of the transcriptional start site (TSS) (**Fig. 3e**). One such BETA was identified within the vicinity of the Otp gene (**Fig. 3f**). This enrichment supports the hypothesis that chromatin accessibility in these cell types is functionally linked to transcriptional activation and that cis-regulatory elements identified from ATAC-seq peaks are predictive of gene expression in a cell-type-specific manner.

### Identifying evolutionarily conserved CREs between mice and humans linked to obesity

Next, to identify mouse cis-regulatory elements (CREs) with the highest likelihood of functional relevance to human biology, we focused on cluster- and cluster-family-specific OCRs that are conserved between mouse and human. To accomplish this, mouse OCR coordinates (mm10) were lifted over to the human genome (hg38) using UCSC’s liftOver tool (UCSC Genome Browser) (**Fig. 4a**) ^56^. With this approach, we identified thousands of orthologous OCRs across each of our clusters. We confirmed that the number of total OCRs for each cluster was not positively correlated with the number of orthologous peaks for that cluster (**Fig. 4b**). These conserved, cell-type-specific OCRs represent strong candidates for encoding transcriptional programs that are both cell-type-specific and functionally conserved across species.

**Figure 4.**
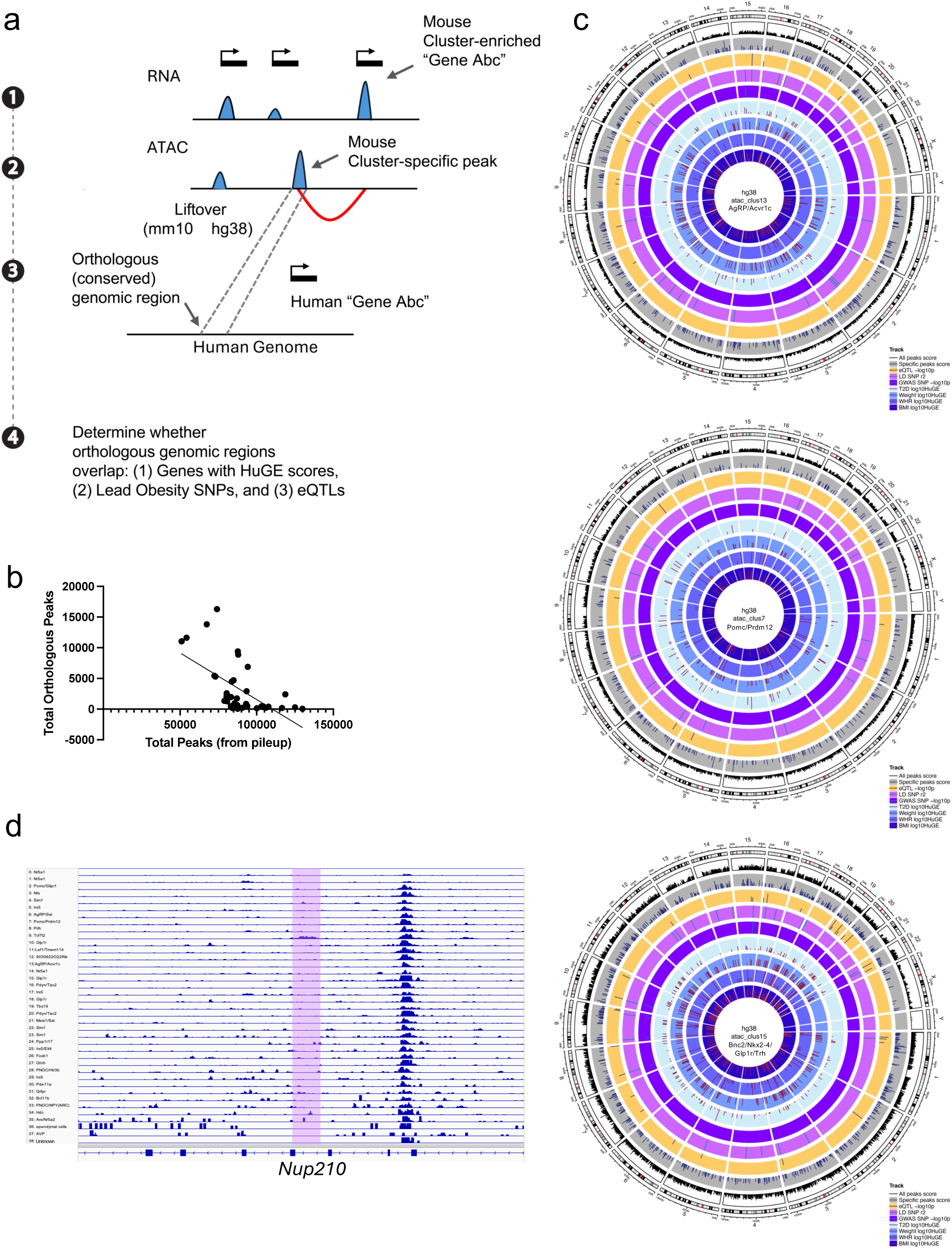
Identifying conserved OCRs associated with obesity-risk variants. **(A) Crude** schematic showing the heuristic used to identify mouse orthologous OCRs that overlap human variants and genes linked to obesity. (B) Relationship between the total number of peaks identified per cluster and the number of orthologous peaks retained for cross-species analysis. (C) Circos plots for clusters 13: Agrp/Sst, 7:Pomc/Prdm12, and 15: Bnc2/Nkx2-4/Glp1r/Trh, each integrating multiple genomic layers. From outer to inner rings, the plots depict: (1) all human homologous OCRs (black), (2) cluster-specific homologous OCRs (blue), (3) genome-wide significant hypothalamus eQTLs (red, −log₁₀p), (4) LD- and (5) Lead-SNP variants associated (p < 1×10^-5^) with BMI, body weight, WHR, and T2D, HuGE-association scores (log₁₀ scale) for (6) type 2 diabetes (cyan), (7) weight (lavender), (8) waist-to-hip ratio (green), and (9) BMI (orange). These integrated views highlight genomic regions where chromatin accessibility, gene regulatory potential, and obesity-related genetic risk converge. (D) snATAC-seq tracks displayed on IGV, showing the Nup210 gene locus, with a lavender overlay highlighting an cluster 9: Tcf7l2-specific OCRs that overlaps a HuGE genes and LD-SNPs.

We then asked whether our cell type–specific orthologous OCRs are proximal (within ±1 Mb) to genes implicated in the control of body weight via human genome-wide association studies (GWAS). Thus, we used the Human Genetic Evidence (HuGE) framework, which synthesizes evidence from large-scale GWAS of common variation, rare-variant association tests, and fine-mapping/colocalization analyses to derive a unified quantitative gene-level score ^57^. Higher scores mean stronger and more consistent human evidence that changes in that gene affect the trait. We annotated the human genes near each orthologous OCR with their HuGE scores for BMI, body weight, waist-to-hip ratio, and Type 2 Diabetes (T2D), and grouped them using the standard tiers (Moderate ≥3, Strong ≥10, Very Strong ≥30, Extreme ≥100, Compelling ≥350).

For each cluster and cluster family, we then counted how many orthologous OCRs sit near genes in these tiers (≥ Stronger) thereby highlighting conserved regulatory neighborhoods where adjacent genes already have substantial human genetic backing for obesity-related phenotypes (**Fig. S4a**).

We next asked whether orthologous OCRs overlap with human GWAS signals for obesity-related phenotypes. Using the NHGRI-EBI GWAS Catalog, we intersected OCRs with variants associated (p < 1×10^-5^) with BMI, body weight, WHR, and T2D by base-pair overlap. To capture correlated, potentially causal variants not reported as leads, we expanded each trait’s lead set with linkage disequilibrium (LD) proxies computed in the 1000 Genomes European panel using PLINK (±1 Mb window, r² ≥ 0.8), and repeated the intersection. For each overlapping OCR, we recorded the strongest GWAS support (maximum -log₁₀p across traits) and the maximum r² of any overlapping proxy, providing a quantitative readout of both association strength and LD proximity to the reported lead. At the cluster level, we report counts of overlaps per trait as well as the distribution of GWAS -log₁₀p and LD r², highlighting conserved OCRs that sit within tightly correlated GWAS neighborhoods for obesity-related traits (**Fig. S4b**).

To evaluate regulatory plausibility in the relevant tissue, we intersected orthologous OCRs with human hypothalamus eQTLs from the EBI eQTL Catalogue (GTEx v8 “Brain_Hypothalamus”), using a discovery threshold of p < 1×10^-5^ and requiring direct base-pair overlap. For overlapping OCRs we retained the rsID(s) and -log₁₀(p), thereby prioritizing conserved, cell type-specific accessible elements that also harbor variants associated with gene expression in human hypothalamus^58^ (**Fig. S4c**). While bulk-tissue eQTLs do not resolve cell types, the intersection with cluster-restricted OCRs points to plausible subtype-specific regulatory variants.

Finally, we integrated these layers visually with Circos plots for each cluster ^59^. From outer to inner rings, the plots display: all human-orthologous OCRs, cluster-specific orthologous OCRs, hypothalamus eQTLs (-log_10_p), LD support (maximum r²), GWAS support (strongest −log_10_p across BMI/weight/WHR/T2D), and log_10_(HuGE) for each trait. These summaries highlight convergent loci where conserved chromatin accessibility, human association signals (direct or high-r² proxies), and tissue-relevant regulatory effects co-localize near genes with strong HuGE support—pinpointing LepR^Hypo^ subtypes and regulatory intervals with the highest translational relevance to human energy balance and obesity risk. Clusters 7, 13, and 15 stood out with enriched overlap between cluster-specific OCRs, significant eQTLs, and high HuGE scores, pointing to specific LepR^Hypo^ subtypes where common functional variants may confer subtype-specific influences on energy balance and obesity risk (**Fig. 4c**). One prime example of a mouse OCRs that is orthologous to a human genomic region with an obesity-associated SNP and eQTL is a 9: Tcf7l2 cluster-specific OCR within the domain of the Nup210 gene (**Fig. 4d**). The Nup210 gene has “Very Strong” Huge Scores of 45 for BMI, T2D and Weight, and the orthologous human genomoic region overlaps LD SNPs associated with BMI and T2D with high r^2^ values.

## DISCUSSION

In this study we enriched LepR-expressing neuronal types prior to generating single-nucleus transcriptional and chromatin accessibility profiles. This approach enabled us to conserve our sequencing reads, while only allocating them towards the goal of profiling the key neuronal types in which we are chiefly interested. As a result we were able to profile these select LepR-expressing neuronal types with unprecedented depth, and consequently managed to identify several novel LepR-expressing neuronal subtypes.

We recapitulated a previous observation of their being two AgRP neuronal subtypes, previously shown to be enriched for *Sst* and *Gm8773*, respectively^10^. We demonstrated that the *AgRP*/*Gm8773* subcluster is greatly enriched for Acvr1c. Moreover, this population (13: AgRP/Acvr1c) is more ventrally located within the ARC than the other (6: AgRP/Sst) which is more dorsally positioned. Moreover, the considerable enrichment of *Acvr1c* suggests Acvr1c signaling within AgRP neuron may be physiologically relevant and potentially impact energy homeostasis. Previous studies, in mice, have shown that Acvr1c loss-of-function mutations are protective against diet-induced obesity, and human SNPs in Acvr1c are associated with reduced body weight ^60, 61^. Whereas the impact of Acvr1c expression and signaling within adipose tissue has been studied, future studies should investigate the role of Acvr1c signaling within the 13: AgRP/Acvr1c neuronal subtype in the control of energy homeostasis.

Several studies have detailed the importance of neuronal types that dually express Lepr and Glp1r within the hypothalamus. LepRb^Glp1r^ neurons have been shown to be critical mediators of Leptin and Glp1r signaling in mice, and evolutionarily conserved in primates and humans ^39^.

Recent studies have also identified Lepr- and Glp1r-expressing neurons within the ARC that suppress food intake when activated, promote feeding when inactivated, and send inhibitory inputs to AgRP neurons, referred to a Trh/Cxcl12 and Bnc2 neurons, respectively ^17, 18^.

Furthermore, LepR^Glp1r^ in the DMH have also been shown to mediate the satiety-inducing effects of the Glp1r agonist Liraglutide and activation of these neurons promotes satiety via their direct inhibition of downstream AgRP neurons^62^. Our findings support the notion that *Bnc2* is a marker of the broad family of Lepr/Glp1r-co-expressing neurons within not only the ARC, but also the DMH, and it is also co-enriched in these neurons with *Nkx2-4*, a previously identified marker for a hypothalamic neuronal types ^41^. Furthermore, we confirm that Trh/Cxcl12 neurons are a major subtype of *Lepr*- and *Glp1r*-expressing neurons, and are discernible from those that are enriched for *Ebf1*, which are predominantly found in the DMH, and those that are enriched for *Tbx19* (but not *Anxa2*), which are predominantly found in the ARC. Whereas *Bnc2* was enriched in each our *Lepr*/*Glp1r*-expressing subtypes (Ebf1, Trh, Txb19), 30% to 43% of *Lepr*/*Glp1r*-expressing neurons, in the ARC and DMH, respectively, did not express Bnc2 according to our Xenium data. It is possible that Bnc2 expression is marginally low and thus dropped out due to a limitation in the sensitivity of our assay, or Bnc2 is simply not expressed uniformly in all *Lepr*/*Glp1r*-expressing neurons in the ARC and DMH. Indeed, 15: Bnc2/Nkx2-4/Glp1r/Trh expressed the most *Lepr*, whereas 10: Bnc2/Nkx2-4/Glp1r/Ebf1 neurons expressed the most *Glp1r* of the three, and 18:Bnc2/Nkx2-4/Glp1r/Tbx19 were relatively most enriched for *Bnc2*. The finding that Bnc2^ARC^ neurons are not responsive Glp1 is surprising^17^, given the strong evidence that this broad family of neurons are enriched for Glp1r, however, given the heterogeneity within the Bnc2 family, it is possible that Glp1 has differing effects on each of the Bnc2 neuron subtypes. If so, an appreciable response owed to Glp1r across all diverse Bnc2 neurons might not be expected. Thus, future experiments should assess the possibility of functional diversity across the Ebf1-, Trh-, and Tbx19-expressing subtypes of Bnc2 neurons in response to exogenous Glp1 and other metabolic perturbations. Finally, given the recent finding that LepR^DMH^ neurons are activated by Liraglutide and mediate its satiety-inducing effects by inhibiting AgRP neurons ^62^, coupling with our finding that *Ebf1*-enriched Bnc2/Nkx2-4/Glp1r/Ebf1 neurons are not only the predominant LepR^Glp1r^ neuronal type within the DMH, but also express the most Glp1r relative to other identified LepR^Glp1r^ neurons (i.e., *Trh*-enriched, and *Tbx19*-enriched), future studies should determine if this *Ebf1*-enriched sub-population is the principal satiety-evoking, Glp1r-responsive, *Lepr*-expressing neuronal population in the DMH. What we’re only slowly beginning to appreciate are the diverse assortment of OCRs that are uniquely found in particular families of cells, and often within particular neuronal subtypes.

These cell-type specific OCRs have the potential to differentially govern the DNA binding substrates of particular transcription factors, dictating when, where, and how particular TFs bind along the genome. Our study reveals a tremendous degree of cell-type specificity of OCRs within the large family of leptin-responsive cell-types within the hypothalamus. We identified thousands of unique OCRs across our cell-types. Again, the existence of cell-type specific OCRs makes it entirely possible that particular non-coding regions exert their effects on particular neuronal types within the hypothalamus. One limitation of our study involves our only characterizing neurons in which LepR-ires-cre is expressed. Non-lepR-expressing neurons have been shown to influence body weight, such as GABAergic non-lepR neurons within the ARC ^63^. Thus, future experiments might consider enriching for non-lepR-expressing neuronal types excluded from our analysis. Furthermore, comparisons between the chromatin profiles revealed in LepR^Hypo^ neurons with those from other cell-types beyond the hypothalamus and across the rodent and human body will be important to identify OCRs in LepR^Hypo^ subtypes that are truly cell-type specific.

Efforts should also be made to uncover masked cell-type specific enhancers that are only accessible in response to particular perturbations. For instance, our previous study involving mouse AgRP neurons identified 2452 fasted-opened, and 203 leptin-opened, areas of chromatin, with a number of these peaks being undetectable prior to fasting, or leptin-treatment, respectively ^7^. Thus, we predict there to be numerous cell-type specific enhancers that are masked in inaccessible regions and waiting to be identified after the proper perturbation.

Furthermore, future studies should characterize the full host of dynamically altered chromatin accessibility changes across various hypothalamic neuronal types in response to perturbations such as a 24-hour, non-torpor inducing, fast, acute and prolonged high-fat diet exposure, and other metabolic perturbations, as way to identify transcriptional pathways that are shared and distinct between different neuronal types.

In this study we were interested in identifying evolutionarily conserved OCRs between mice and humans to: (1) identify OCRs that seem to be quite important, in regulating the expression of genes that influence energy homeostasis across species, (2) identify enhancers that when perturbed can disrupt gene expression across species, (3) hone in on those coding and non-coding variants with the potential to influence gene expression within discrete neuronal subtypes, while ultimately increasing or decreasing the risk for developing obesity.

We identified a rich dataset of tens of Lead SNPs, and hundreds of LD-SNP, associated with body weight, and hundreds of eQTL functional variants, that overlapped orthologous OCRs found across our mouse LepR^Hypo^ neuronal types. A recent paper showed that a major obesity-associated SNP, expressed in mice, increased body weight by influencing Irx3-expressing neurons, thereby showcasing how a specific SNP can mechanistically increase the risk for developing obesity ^64^. Thus, future experiments should determine whether particular SNPs and eQTLs overlapping cell-type specific orthologous enhancers (OCRs) (1) influence gene expression in various *in vitro* and *in vivo* models, and (2) influence energy homeostasis, and, in particular, susceptibility to diet-induced obesity.

## LEAD CONTACT AND MATERIALS AVAILABILITY

Further information and requests for resources and reagents should be directed to and will be fulfilled by the lead contact, Frankie D. Heyward.

## EXPERIMENTAL MODEL AND SUBJECT DETAILS

### Mouse Models

#### Generation of mice

We crossed transgenic Nuclear tagging and Translating Ribosome Affinity Purification (NuTRAP) mice with Lepr-IRES-Cre (JAX #: 032457) mice to generate the NuTRAP^Lepr^ mouse line, from which we could isolate LepR neuron-specific mRNA and nuclei. Mice maintained on a C57Bl/6J background, were used for all studies.

#### Animals: standard fed, fasted, and leptin-treated comparison

Animal experiments for snRNA-seq/snATAC-seq were performed with approval from the Institutional Animal Care and Use Committees of The Harvard Center for Comparative Medicine and Beth Israel Deaconess Medical Center. 1 year old female C57BL/6J NuTRAP^Lepr^ mice were fed a standard chow diet *ad libitum*.

#### Isolation of LepR neuronal nuclei from NuTRAP Mice

LepR neuronal nuclei were isolated as previously described, with minor alterations. Using a fluorescent stereoscope, dissected whole hypothalamus from 1-old female mice were collected, snap frozen, and stored at −80C. 6 isolated hypothalami were pooled, dounce homogenized in nuclear preparation buffer (NPB; 10 mM HEPES [pH 7.5], 1.5mM MgCl_2_, 10 mM KCl, 250 mM sucrose, 0.1% NP-40, and 0.2 mM DTT), and the homogenate was filtered through a 100 μM strainer and centrifuged to pellet the nuclei. Nuclei were washed with NPB, re-suspended in nuclear sorting buffer (10 mM Tris [pH 7.5], 40 mM NaCl, 90 mM KCl, 2 mM EDTA, 0.5 mM EGTA, 0.1% NP-40, 0.2 mM DTT), and filtered again through a 40 μM strainer. Isolated nuclei were sorted by flow cytometry based on LepR neuron-specific GFP expression, and collected into 500-750uL PBS (0.1% NP40, with protease and RNase inhibitors) in 1.5 mL microcentrifuge tubes and stored on ice. With this approach we routinely obtain ∼20,000-40,000 LepR neuronal nuclei per mouse, for a total of ∼120,000 nuclei in all. The tube was spun at 1000 rpm for 10 mins, at 4°C. Supernatant was removed via gentle decantation and nuclei were resuspended in 100 μl of 10x Genomics Diluted Nuclei Buffer.

#### Single-nucleus Multiome library preparation and sequencing

Approximately 5,000 nuclei per library were loaded into individual lanes of the Chromium Next GEM Chip J and processed on the Chromium Controller (10x Genomics, Pleasanton, CA, USA). Transposition and cDNA synthesis, followed by library construction, were carried out according to the manufacturer’s protocol (Chromium Next GEM Single Cell Multiome ATAC + Gene Expression User Guide, Rev. F). Final libraries were normalized, pooled, and sequenced on a NovaSeq SP-100 flow cell (Illumina, San Diego, CA, USA).

#### Demultiplexing and alignment

Sequencing output (BCL files) was demultiplexed into FASTQ files using bcl2fastq (Illumina) and cellranger-arc mkfastq (v2.0.0, 10x Genomics). Four libraries were generated from LepR-expressing hypothalamic nuclei and processed individually with cellranger-arc count using the mouse reference genome mm10 (refdata-cellranger-arc-mm10-2020-A-2.0.0). Libraries were then aggregated with cellranger-arc aggr without normalization to produce a combined dataset.

#### Data import and Seurat object construction

RNA UMI counts and ATAC fragment counts were imported into R and combined in a Seurat object (v4.0) for downstream analysis. Gene annotations were obtained from Ensembl release 98 using AnnotationHub, and ATAC fragments were associated with peak calls to generate a chromatin accessibility assay. Metadata were added to label the four libraries.

#### Quality control

Per-cell quality control metrics were computed using both RNA and ATAC data. Metrics included the number of ATAC transposition events in peaks (nCount_ATAC), number of accessible peaks (nFeature_ATAC), transcription start site (TSS) enrichment score, nucleosome signal, number of RNA UMIs (nCount_RNA), number of expressed genes (nFeature_RNA), and percentage of mitochondrial RNA UMIs (percent_mt). Cells were excluded if they met any of the following thresholds: >100,000 ATAC counts, <100 ATAC counts, <50 peaks, nucleosome signal >2, TSS enrichment <2, >100,000 RNA UMIs, <500 RNA UMIs, <500 expressed genes, or >10% mitochondrial RNA content. After filtering, high-quality nuclei were retained for all downstream analyses.

#### Peak calling

To generate a high-confidence set of chromatin accessibility peaks, we called peaks with MACS2 (Zhang et al., 2008) on the aggregated ATAC fragments. Peaks on nonstandard chromosomes and those overlapping the ENCODE mm10 blacklist regions were removed. A chromatin accessibility assay was then created using this MACS2 peak set and added to the Seurat object for downstream analyses.

#### Gene expression data processing

Gene expression counts were normalized using the SCTransform method (Hafemeister and Satija, 2019). Principal component analysis (PCA) was performed on the top 3,000 variable genes. The number of principal components to retain was determined by Horn’s parallel analysis (Horn, 1965), which supported the use of 53 components for downstream clustering.

#### DNA accessibility data processing

Chromatin accessibility data were processed by latent semantic indexing (LSI) (Stuart et al., 2021). We retained the top features per cell, applied term frequency–inverse document frequency (TF–IDF) normalization, and performed singular value decomposition (SVD) to generate 100 LSI components.

#### Joint integration and clustering

To integrate modalities, we used the weighted nearest neighbor (WNN) method in Seurat (Hao et al., 2021), combining the PCA results from RNA with the LSI results from ATAC (excluding the first LSI dimension, which is correlated with sequencing depth). A joint neighbor graph was computed and clustered using the Louvain algorithm (Blondel et al., 2008). This analysis identified 39 distinct clusters.

#### Visualization

We computed a joint UMAP embedding based on the WNN graph and visualized clusters across all samples (cell_clusters_umap_all.pdf) as well as sample-specific projections (cell_clusters_umap.pdf). Independent UMAP projections were also generated for RNA (PCA-derived) and ATAC (LSI-derived) modalities.

#### Label transfer

Cell type annotation of the LepR-expressing hypothalamic single-nucleus multiome dataset was performed by label transfer using the HypoMap reference atlas. The reference Seurat object, including curated class annotations (Author_Class_Curated, 2-class, 7-class, 25-class, 66-class, predicted region, and summarized region), was downloaded from the University of Cambridge repository. Both reference and query datasets were normalized using LogNormalize, 3,000 variable features were selected, and the data were scaled prior to principal component analysis (30 components). Transfer anchors were identified with the Seurat function FindTransferAnchors using the reference reduction set to PCA, and labels were transferred with TransferData across the same 30 dimensions. Prediction scores were retained for all cells.

Scores greater than 50 were considered reliable assignments, while scores between 33 and 50 were classified as possible. For each cluster, the proportion of predicted labels was tabulated, and the most abundant label was used as the reassigned identity. UMAPs were generated to visualize the transferred annotations across all levels.

#### Differential gene expression analysis

Cluster-specific marker genes were identified using the Wilcoxon rank sum test (Mann and Whitney, 1947). Differentially expressed genes were required to have a log₂ fold-change >0.25 and expression in at least 10% of cells within the cluster. P values were adjusted for multiple testing using Bonferroni correction.

#### Identification of cluster-specific chromatin accessibility peaks

To identify peaks enriched in specific cell clusters, we first generated pseudobulk ATAC-seq profiles by aggregating reads from each cluster using sinto (Stuart et al., 2021). Pseudobulk BAM files were merged across the four libraries with samtools (Li et al., 2009). Peaks were then called with MACS2 (Zhang et al., 2008) using paired-end alignment mode (BAMPE), a false discovery rate (q-value) cutoff of 0.05, and signal per million reads (SPMR) normalization.

Peaks with normalized signal <1 were excluded. Cluster-specific peaks were defined as those present in one cluster but absent from all other clusters. We also identified peaks unique to “cluster families” (sets of closely related clusters) by overlapping and merging peaks within each family and excluding peaks shared with other clusters. Peaks located more than 1 Mb from annotated genes were filtered out.

Remaining peaks were annotated to nearby genes using ChIPseeker and related packages, with genomic features derived from Ensembl (release 98, mm10). Annotation included peak coordinates, enrichment scores, distance to transcription start site, and associated gene identifiers (ENSEMBL ID, Entrez ID, gene symbol, biotype, and name). Expression status of linked genes was determined from pseudobulk RNA-seq counts, with genes considered expressed if counts per million (CPM) exceeded 1.

#### Cross-species mapping of cluster-specific peaks

Mouse cluster- and cluster-family–specific ATAC peaks linked to expressed genes (CPM > 1) were lifted over from mm10 to hg38 using UCSC chain files. Lifted-over human intervals were merged when separated by <100 bp to account for fragmented mappings. Regions >1 Mb from any annotated gene were excluded. Human intervals were annotated to nearest genes and genomic features using TxDb.Hsapiens.UCSC.hg38.knownGene with R/Bioconductor package org.Hs.eg.db.

#### Association with metabolic traits (HuGE)

Human genes linked to lifted-over regions were cross-referenced against the Common Metabolic Diseases Knowledge Portal for type 2 diabetes (T2D) and related phenotypes (BMI, body weight, waist-to-hip ratio). Human Genetic Evidence (HuGE) scores were appended to the gene annotations.

#### Overlap with human hypothalamus eQTLs

eQTLs for human brain hypothalamus (GTEx V8 via the EBI eQTL Catalogue) were intersected with lifted-over regions. Variants with p < 1×10^-^⁵ were retained, and intersecting variants (rsID and p-value) were recorded for each region.

#### Overlap with GWAS loci and LD proxies

Genome-wide significant SNPs for T2D, BMI, body weight, and WHR (EBI GWAS Catalog; threshold p < 1×10⁻⁵) were intersected with lifted-over regions. To capture proxies in linkage disequilibrium, LD was computed with PLINK using the 1000 Genomes EUR reference (window = 1,000 kb; r² ≥ 0.8). For each region, overlapping GWAS SNPs (rsID, p-value) and LD proxies (rsID, r², lead SNP) were summarized.

#### Xenium in situ transcriptomics

Fresh frozen mouse hypothalamic tissue was processed on the 10x Genomics Xenium In Situ platform. Guided by our single-cell Multiome atlas, we designed a custom 300-gene Xenium panel enriched for cluster-specific marker genes to support cell type recognition and subtype analyses. RNA integrity, assessed on adjacent material, was high and exceeded vendor-recommended thresholds. Sections were prepared according to the Xenium In Situ Fresh Frozen Tissue Preparation Guide (10x Genomics), placed in the designated sample area on Xenium slides, and fixed/permeabilized per manufacturer recommendations to facilitate mRNA accessibility. Probe hybridization was performed overnight at 50 °C, unbound probes were removed, and target-bound probes were ligated at 37 °C (Xenium ligase A/B). Circularized probes underwent rolling circle amplification to generate barcoded amplicons; background fluorescence was chemically quenched. Slides were imaged on a Xenium Analyzer for decoding and image acquisition. DAPI nuclear images were used for cell assignment, and built-in machine-learning algorithms performed cell segmentation. Transcript counts per cell were obtained directly from Xenium Explorer using the Cell Selections workflow, with no external segmentation.

For AgRP subtype analyses within the arcuate nucleus (ARC), AgRP neurons were manually selected in a subtype-blind manner using total Xenium transcript counts per cell. Cells with ≥13 total transcripts were included (mean 77; maximum 228), yielding n = 210 cells. Acvr1c and Sst transcript layers were overlaid and per-cell counts extracted; genes were considered detected if ≥2 transcripts were present. Fractional expression was calculated as Acvr1c/(Acvr1c+Sst) and, separately, Sst/(Acvr1c+Sst). Cells were classified as Acvr1c-positive when Acvr1c/(Acvr1c+Sst) > 0.66 (with Acvr1c ≥ 2 transcripts) or Sst-positive when Sst/(Acvr1c+Sst) > 0.66 (with Sst ≥ 2 transcripts). Cells with both genes detected but neither fraction > 0.66 were considered co-expressing, and cells with neither gene detected were considered negative for both.

For LepR/Glp1r analyses in the ARC/VMH and DMH, double-positive cells were first identified using a detection threshold of ≥3 transcripts for both Lepr and Glp1r (ARC/VMH: n = 99; DMH: n = 109). Within these double-positive sets, Bnc2, Ebf1, Trh, and Tbx19 transcript layers were overlaid and per-cell counts extracted. Positivity thresholds were set a priori as follows: Bnc2, ≥1 transcript; Ebf1, Trh, and Tbx19, ≥2 transcripts. For subtype designation among Ebf1/Trh/Tbx19, fractional dominance was computed as gene count/(Ebf1+Trh+Tbx19); cells were labeled Ebf1-, Trh-, or Tbx19-positive if the corresponding fraction exceeded 0.66. In the ARC, Lepr⁺/Glp1r⁺ cells were further stratified by Bnc2 status (Bnc2⁺ vs. Bnc2⁻), and the numbers and proportions of Ebf1-, Trh-, Tbx19-positive, or “Neither” cells were determined within each stratum. All detection and fractional-dominance thresholds were defined a priori to minimize spurious single-transcript calls and to provide conservative subtype assignments from per-cell Xenium counts.

## ACKNOWLEDGEMENTS

The authors would like to thank the Single Cell Core at Harvard Medical School, Boston, MA, for performing the multiome snRNA-seq/snATAC-seq sample preparation. We thank the BIDMC Flow Cytometry Core. We would also like to thank the Bioinformatics and Biostatistics Core at Joslin Diabetes Center for assistance with bioinformatic analyses. We also thank the UT Southwestern Medical Center Microarray and Immune Phenotyping Core, and UT Southwestern Medical Center Histo Pathology Core for their work related to the Xenium sample prep and data acquisition.

## AUTHOR CONTRIBUTIONS

F.D.H. conceived this study and interpreted the results of all experiments. F.D.H supervised this study. F.D.H conceived of the cluster- and cluster-family specific peak enrichment analysis, mouse snATAC-seq integration with human obesity SNP and eQTL results, and Circos plot visualization. H.P. performed computational and bioinformatic analyses involving sequencing data and was supervised by J.M.D. F.D.H wrote the manuscript with added input from H.P. and J.M.D.

## DECLARATION OF INTERESTS

None.

---

**Supplementary Figure 1.**
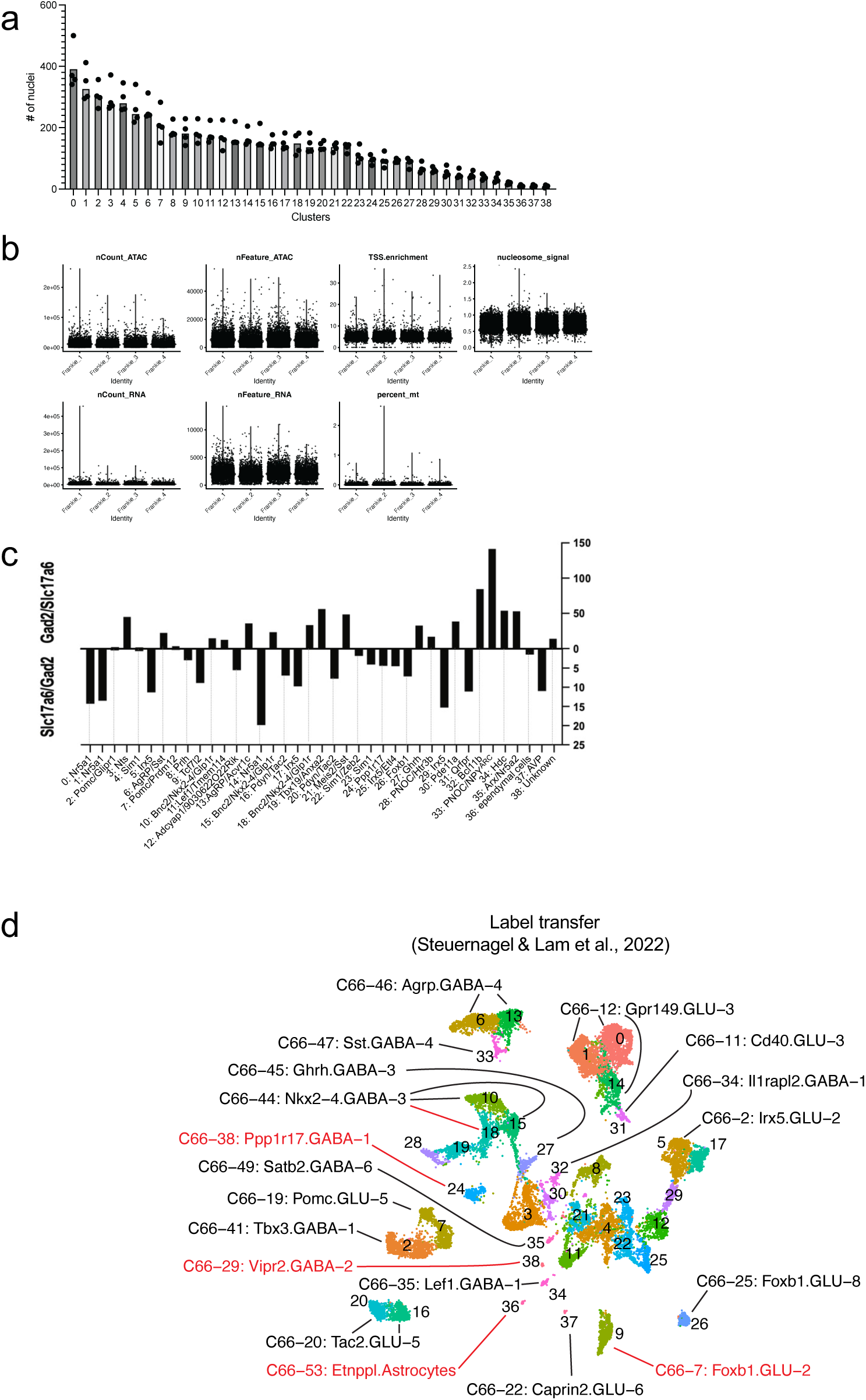
snRNA-seq quality control summary statistics. Associated with main Figure 1. (A) Number of nuclei for all 4 libraries, across all 39 clusters. (B) Quality control metrics for snRNA-seq and snATAC-seq libraries. Distribution of ATAC fragment counts (nCount_ATAC), detected accessible features (nFeature_ATAC), transcription start site enrichment (TSS.enrichment), nucleosome signal, RNA transcript counts (nCount_RNA), detected genes (nFeature_RNA), and mitochondrial transcript percentage (percent_mt) across all four libraries. (C) Ratio of glutamateric marker (Slc17a6) to GABAergic marker (Gad2) across our clusters.

**Supplementary Figure 2.**
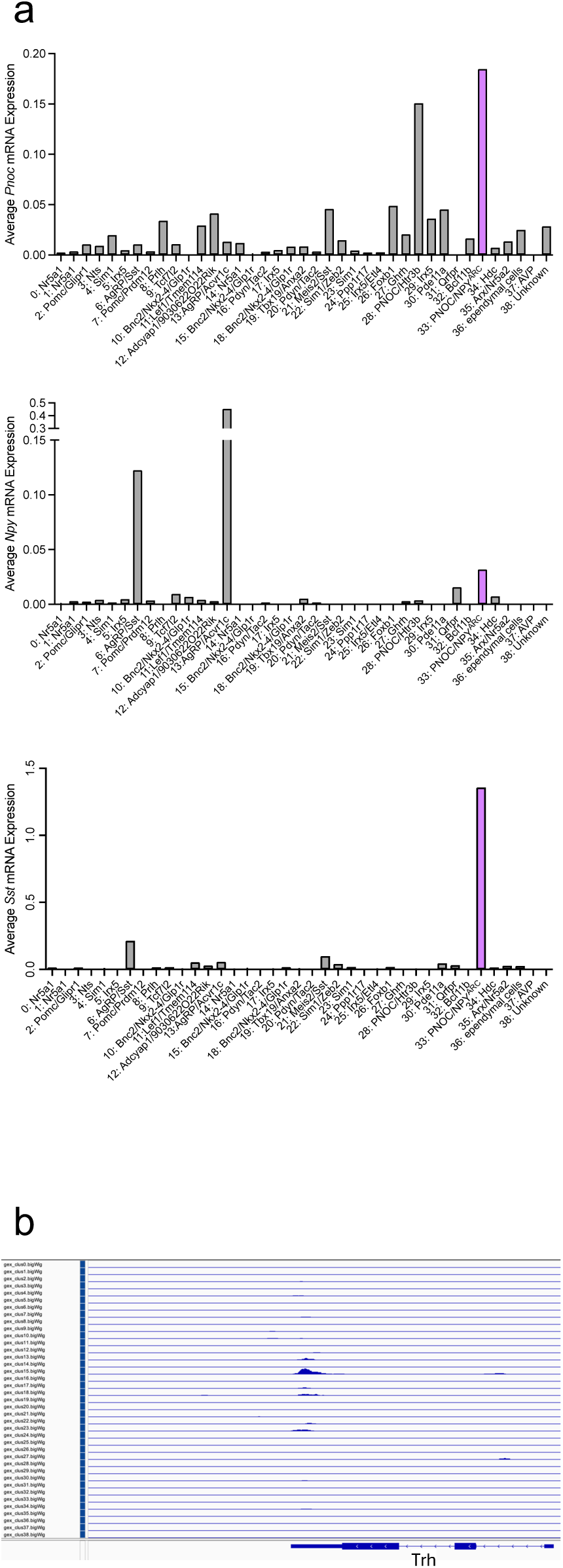
Gene-expression profiles within neuronal subtypes. Associated with main Figure 2. (A) Pnoc, Npy, and Sst expression across clusters. (B) snRNA-seq tracks displayed in IGV showing the Trh gene locus.

**Supplementary Figure 3.**
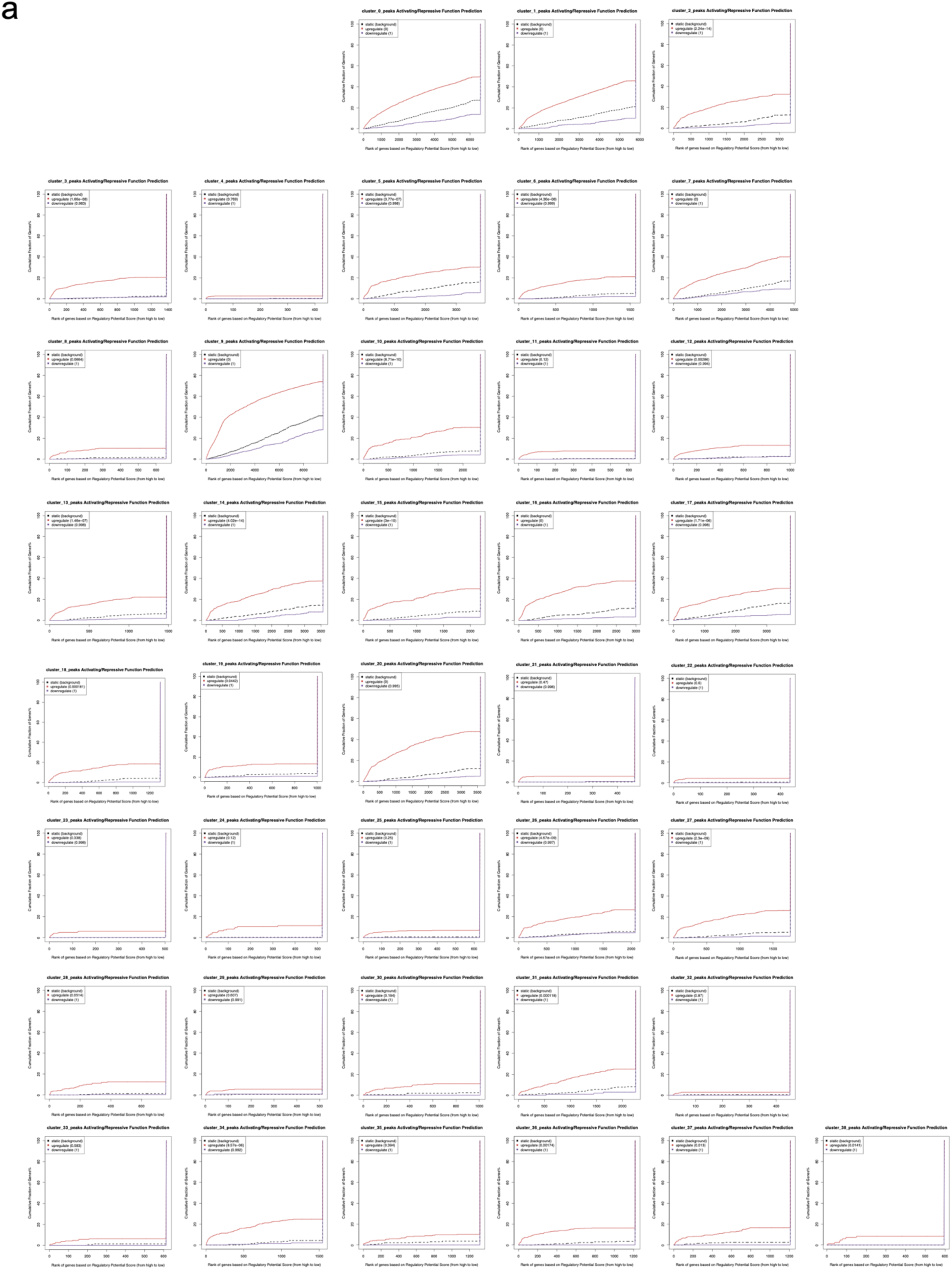
Functional prediction of cluster-specific snATAC-seq peaks using BETA. Associated with main. Figure 3. (A) Cumulative distribution function (CDF) plots from BETA analysis showing the predicted activating (red) or repressive (black) regulatory potential of cluster-specific accessible regions on gene expression, with significant deviations of the red (upregulated) or black (downregulated) curves from the background (dashed line) indicating predicted transcriptional activating or repressive functions, for each of our 39 clusters.

**Supplementary Figure 4.**
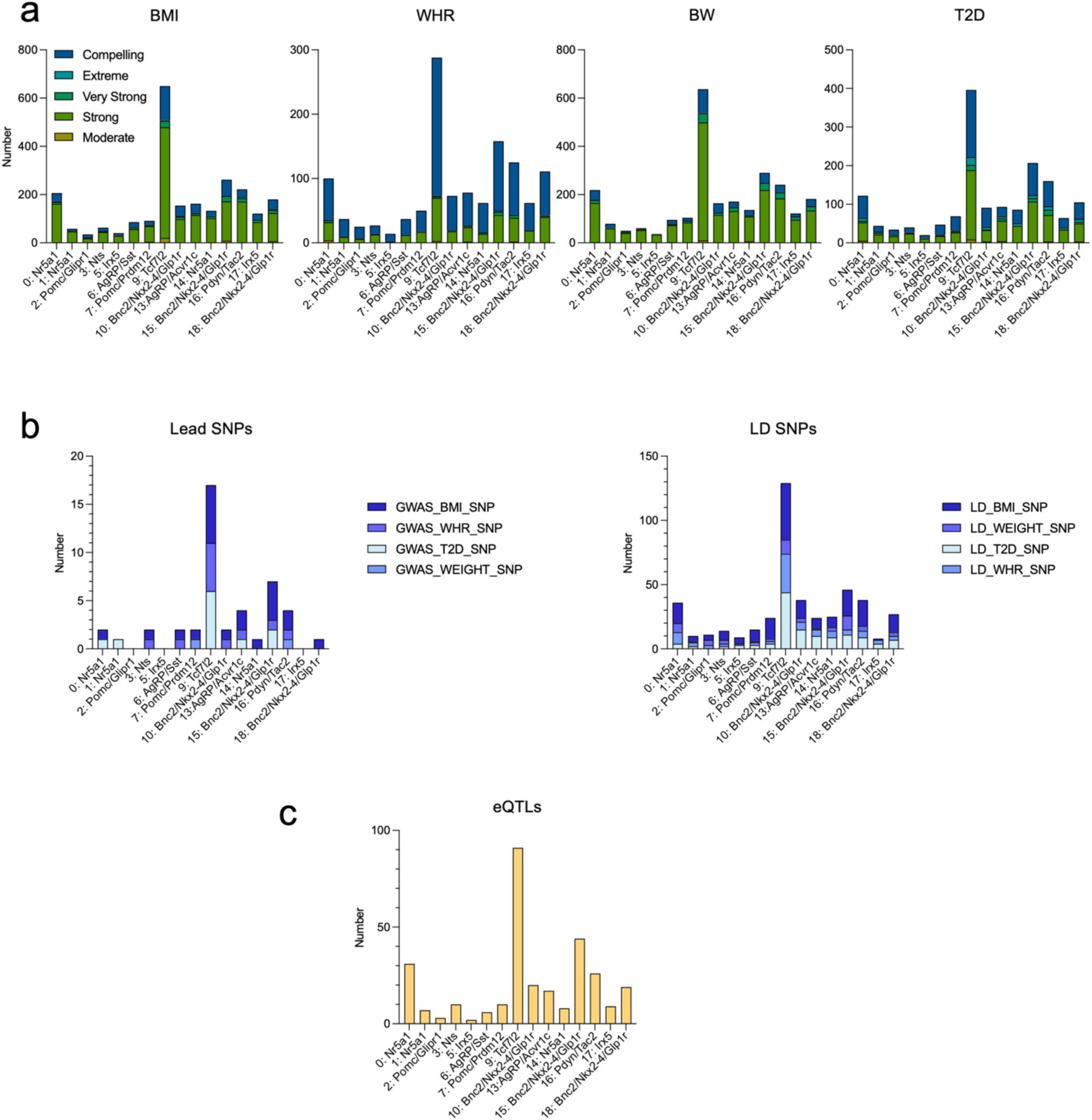
Integration of HuGE scores, GWAS SNPs, and eQTLs with hypothalamic OCR clusters. Associated with main. Figure 4. (A) Distribution of orthologous OCR–associated genes across Human Genetic Evidence (HuGE) score tiers (Moderate, Strong, Very Strong, Extreme, Compelling) for body mass index (BMI), waist-to-hip ratio (WHR), body weight (BW), and type 2 diabetes (T2D). (B) Overlap of cluster-specific OCRs with genome-wide association study (GWAS) lead SNPs (left) and SNPs in linkage disequilibrium (LD) with GWAS loci (right) across the same traits. (C) Number of expression quantitative trait loci (eQTLs) overlapping OCRs, highlighting putative regulatory variants linked to gene expression in particular LepR^Hypo^ subtypes.

## Notes

### Competing Interest Statement

The authors have declared no competing interest.

### Funding

- Simons Foundation: SFI-AN-NC-AB-Independence Faculty-00006766
- National Institutes of Health: K01 DK131347

## References

1. Ahima RS, et al. Role of leptin in the neuroendocrine response to fasting. Nature 382, 250–252 (1996).

2. Friedman JM. Leptin and the endocrine control of energy balance. Nat Metab 1, 754–764 (2019).

3. Zhang Y, Proenca R, Maffei M, Barone M, Leopold L, Friedman JM. Positional cloning of the mouse obese gene and its human homologue. Nature 372, 425–432 (1994).

4. Chen H, et al. Evidence that the diabetes gene encodes the leptin receptor: identification of a mutation in the leptin receptor gene in db/db mice. Cell 84, 491–495 (1996).

5. Xu J, et al. Genetic identification of leptin neural circuits in energy and glucose homeostases. Nature 556, 505–509 (2018).

6. Allison MB, Patterson CM, Krashes MJ, Lowell BB, Myers MG, Jr., Olson DP. TRAP-seq defines markers for novel populations of hypothalamic and brainstem LepRb neurons. Mol Metab 4, 299–309 (2015).

7. Heyward FD, et al. AgRP neuron cis-regulatory analysis across hunger states reveals that IRF3 mediates leptin’s acute effects. Nat Commun 15, 4646 (2024).

8. Cedernaes J, et al. Transcriptional Basis for Rhythmic Control of Hunger and Metabolism within the AgRP Neuron. Cell Metab 29, 1078–1091 e1075 (2019).

9. Tang Q, et al. Leptin receptor neurons in the dorsomedial hypothalamus input to the circadian feeding network. Sci Adv 9, eadh9570 (2023).

10. Campbell JN, et al. A molecular census of arcuate hypothalamus and median eminence cell types. Nature Neuroscience 2017 20:*3* 20, (2017-02-06).

11. Chen R, Wu X, Jiang L, Zhang Y. Single-Cell RNA-Seq Reveals Hypothalamic Cell Diversity. Cell Reports 18, (2017/03/28).

12. Kim DW, et al. The cellular and molecular landscape of hypothalamic patterning and differentiation from embryonic to late postnatal development. Nat Commun 11, 4360 (2020).

13. Yu H, Rubinstein M, Low MJ. Developmental single-cell transcriptomics of hypothalamic POMC neurons reveal the genetic trajectories of multiple neuropeptidergic phenotypes. Elife 11, (2022).

14. Mickelsen LE, et al. Single-cell transcriptomic analysis of the lateral hypothalamic area reveals molecularly distinct populations of inhibitory and excitatory neurons. Nature Neuroscience 2019 22:*4* 22, (2019-03-11).

15. Hajdarovic KH, et al. Single-cell analysis of the aging female mouse hypothalamus. Nat Aging 2, 662–678 (2022).

16. Junaid M, Choe HK, Kondoh K, Lee EJ, Lim SB. Unveiling Hypothalamic Molecular Signatures via Retrograde Viral Tracing and Single-Cell Transcriptomics. Sci Data 10, 861 (2023).

17. Tan HL, et al. Leptin-activated hypothalamic BNC2 neurons acutely suppress food intake. Nature 636, 198–205 (2024).

18. Webster AN, et al. Molecular connectomics reveals a glucagon-like peptide 1-sensitive neural circuit for satiety. Nat Metab 6, 2354–2373 (2024).

19. Griffith EC, West AE, Greenberg ME. Neuronal enhancers fine-tune adaptive circuit plasticity. Neuron 112, 3043–3057 (2024).

20. Friedman MJ, Wagner T, Lee H, Rosenfeld MG, Oh S. Enhancer-promoter specificity in gene transcription: molecular mechanisms and disease associations. Exp Mol Med 56, 772–787 (2024).

21. Inoue F, et al. Genomic and epigenomic mapping of leptin-responsive neuronal populations involved in body weight regulation. Nature Metabolism 2019 1:*4* 1, (2019-04-08).

22. Buenrostro JD, et al. Single-cell chromatin accessibility reveals principles of regulatory variation. Nature 523, 486–490 (2015).

23. Ma S, et al. Chromatin Potential Identified by Shared Single-Cell Profiling of RNA and Chromatin. Cell 183, 1103–1116 e1120 (2020).

24. Luo C, et al. Single nucleus multi-omics identifies human cortical cell regulatory genome diversity. Cell Genom 2, (2022).

25. Kim DW, et al. Decoding gene networks controlling hypothalamic and prethalamic neuron development. Cell Reports 44, (2025/06/24).

26. Ferris E, et al. Genomic convergence in hibernating mammals elucidates the genetics of metabolic regulation in the hypothalamus. Science 389, (2025-07-31).

27. Leshan RL, Bjornholm M, Munzberg H, Myers MG, Jr. Leptin receptor signaling and action in the central nervous system. Obesity (Silver Spring) 14 **Suppl 5**, 208S–212S (2006).

28. Roh HC, Tsai LT, Lyubetskaya A, Tenen D, Kumari M, Rosen ED. Simultaneous Transcriptional and Epigenomic Profiling from Specific Cell Types within Heterogeneous Tissues In Vivo. Cell Rep 18, 1048-1061 (2017).

29. Dhillon H, et al. Leptin Directly Activates SF1 Neurons in the VMH, and This Action by Leptin Is Required for Normal Body-Weight Homeostasis. Neuron 49, (2006/01/19).

30. Kim KW, Sohn J-W, Kohno D, Xu Y, Williams K, Elmquist JK. SF-1 in the ventral medial hypothalamic nucleus: A key regulator of homeostasis. Molecular and Cellular Endocrinology 336, (2011/04/10).

31. Biglari N, et al. Functionally distinct POMC-expressing neuron subpopulations in hypothalamus revealed by intersectional targeting. Nat Neurosci 24, 913–929 (2021).

32. Leon S, et al. Single cell tracing of Pomc neurons reveals recruitment of ‘Ghost’ subtypes with atypical identity in a mouse model of obesity. Nat Commun 15, 3443 (2024).

33. Woodworth HL, et al. Lateral Hypothalamic Neurotensin Neurons Orchestrate Dual Weight Loss Behaviors via Distinct Mechanisms. Cell Rep 21, 3116–3128 (2017).

34. Brown JA, Wright A, Bugescu R, Christensen L, Olson DP, Leinninger GM. Distinct Subsets of Lateral Hypothalamic Neurotensin Neurons are Activated by Leptin or Dehydration. Sci Rep 9, 1873 (2019).

35. Naganuma F, Kroeger D, Bandaru SS, Absi G, Madara JC, Vetrivelan R. Lateral hypothalamic neurotensin neurons promote arousal and hyperthermia. PLoS Biol 17, e3000172 (2019).

36. Son JE, Dou Z, Kim KH, Hui CC. Deficiency of Irx5 protects mice from obesity and associated metabolic abnormalities. Int J Obes (Lond) 46, 2029–2039 (2022).

37. Son JE, et al. Irx3 and Irx5 in Ins2-Cre(+) cells regulate hypothalamic postnatal neurogenesis and leptin response. Nat Metab 3, 701–713 (2021).

38. Campbell JN, et al. A molecular census of arcuate hypothalamus and median eminence cell types. Nature Neuroscience 2017 20:*3* 20, (2017-02-06).

39. Rupp AC, et al. Suppression of food intake by Glp1r/Lepr-coexpressing neurons prevents obesity in mouse models. J Clin Invest 133, (2023).

40. Lee S, Lee CE, Elias CF, Elmquist JK. Expression of the diabetes-associated gene TCF7L2 in adult mouse brain. Journal of Comparative Neurology 517, (2009/12/20).

41. Steuernagel L, et al. HypoMap—a unified single-cell gene expression atlas of the murine hypothalamus. Nature Metabolism 2022 4:*10* 4, (2022-10-20).

42. Tadross JA, et al. A comprehensive spatio-cellular map of the human hypothalamus. Nature 2025 639:8055 639, (2025-02-05).

43. Krashes MJ, et al. An excitatory paraventricular nucleus to AgRP neuron circuit that drives hunger. Nature 507, 238–242 (2014).

44. Moore AM, Coolen LM, Porter DT, Goodman RL, Lehman MN. KNDy Cells Revisited. Endocrinology 159, (2018/09/01).

45. Caglar C, Friedman J. Restriction of food intake by PPP1R17-expressing neurons in the DMH. Proc Natl Acad Sci U S A 118, (2021).

46. Bilella A, Alvarez-Bolado G, Celio MR. The Foxb1-expressing neurons of the ventrolateral hypothalamic parvafox nucleus project to defensive circuits. Journal of Comparative Neurology 524, (2016/10/15).

47. Lee B, et al. Dlx1/2 and Otp coordinate the production of hypothalamic GHRH- and AgRP-neurons. Nature Communications 2018 9:*1* 9, (2018-05-23).

48. Mani BK, et al. The role of ghrelin-responsive mediobasal hypothalamic neurons in mediating feeding responses to fasting. Mol Metab 6, 882–896 (2017).

49. Solheim MH, et al. Hypothalamic PNOC/NPY neurons constitute mediators of leptin-controlled energy homeostasis. Cell 188, (2025/06/26).

50. Affinati AH, et al. Cross-species analysis defines the conservation of anatomically segregated VMH neuron populations. Elife 10, (2021).

51. Solheim MH, et al. Hypothalamic PNOC/NPY neurons constitute mediators of leptin-controlled energy homeostasis. Cell 188, 3550–3566 e3522 (2025).

52. Fujita A, et al. Hypothalamic Tuberomammillary Nucleus Neurons: Electrophysiological Diversity and Essential Role in Arousal Stability. The Journal of Neuroscience 37, (2017 Sep 27).

53. Pei H, Sutton AK, Burnett KH, Fuller PM, Olson DP. AVP neurons in the paraventricular nucleus of the hypothalamus regulate feeding. Molecular Metabolism 3, (2014 Jan 8).

54. Hentges ST, Otero-Corchon V, Pennock RL, King CM, Low MJ. Proopiomelanocortin expression in both GABA and glutamate neurons. J Neurosci 29, 13684–13690 (2009).

55. Wang S, et al. Target analysis by integration of transcriptome and ChIP-seq data with BETA. Nat Protoc 8, 2502–2515 (2013).

56. Perez G, et al. The UCSC Genome Browser database: 2025 update. Nucleic Acids Res 53, D1243–D1249 (2025).

57. Dornbos P, et al. Evaluating human genetic support for hypothesized metabolic disease genes. Cell Metab 34, 661–666 (2022).

58. Kerimov N, et al. A compendium of uniformly processed human gene expression and splicing quantitative trait loci. Nat Genet 53, 1290–1299 (2021).

59. Krzywinski M, et al. Circos: An information aesthetic for comparative genomics. Genome Research 19, (2009-09-01).

60. Emdin CA, et al. DNA Sequence Variation in ACVR1C Encoding the Activin Receptor-Like Kinase 7 Influences Body Fat Distribution and Protects Against Type 2 Diabetes. Diabetes 68, 226–234 (2019).

61. Tangseefa P, Jin H, Zhang H, Xie M, Ibanez CF. Human ACVR1C missense variants that correlate with altered body fat distribution produce metabolic alterations of graded severity in knock-in mutant mice. Mol Metab 81, 101890 (2024).

62. Kim KS, et al. GLP-1 increases preingestive satiation via hypothalamic circuits in mice and humans. Science 385, 438–446 (2024).

63. Li H, Su C, Xu Y, Ludwig MQ, Davis J, Tong Q. An alternative neural basis underlying leptin resistance. Cell Rep 44, 115863 (2025).

64. Sullivan AI, et al. Mice harboring the obesity-associated SNP rs1421085 exhibit increased body weight and reveal an IRX3 neuronal circuit regulating body weight. Mol Metab 100, 102234 (2025).

